# Hypoxia drives HIF2-dependent macrophage cell cycle entry and susceptibility to lentiviral transduction

**DOI:** 10.1101/2023.12.15.571908

**Authors:** Bo Meng, Na Zhao, Petra Mlcochova, Isabella A.T.M Ferreira, Brian M. Ortmann, Tanja Davis, Niek Wit, Jan Rehwinkel, Simon Cook, Patrick H. Maxwell, James A. Nathan, Ravindra K. Gupta

## Abstract

Macrophages play critical roles across health and disease. Low oxygen conditions (hypoxia) have been associated primarily with cell cycle arrest in cultured dividing cells. Macrophages are typically quiescent in G0, though yolk sac and bone marrow derived macrophages frequently proliferate and monocyte-derived tissue macrophages are able to proliferate in response to tissue signals. Here we show that hypoxia (1% oxygen tension) results in reversible entry into the cell cycle in monocyte derived macrophages (MDM) and mouse peritoneal macrophages. Cell cycle progression is largely limited to G1/S phase with very little progression to G2/M. Mechanistically, this cell cycle transitioning is triggered by a HIF2α-directed transcriptional program. The response is accompanied by increased expression of cell cycle-associated proteins, including CDK1, and reversible activation of the canonical mitogen-activated MEK-ERK proliferation pathway. CDK1 associated SAMHD1 phosphorylation at T592 in hypoxic macrophages renders them hyper-susceptible to lentiviral transduction. Furthermore, PHD inhibitors, which activate HIFs, are able to recapitulate HIF2α-dependent cell cycle entry in macrophages, as well as susceptibility to lentiviral transduction. Finally, we demonstrate that tumour associated macrophages (TAM) in lung cancers exhibit transcriptomic profiles representing responses to low oxygen and cell cycle progression at single cell level. This work uncovers HIF2α driven macrophage cell cycle progression in low oxygen conditions that culminates in SAMHD1 phosphorylation and high susceptibility to lentiviral transduction. These findings have implications for inflammation, neoplasia and pathogen defence.

## Introduction

Macrophages are typically quiescent and reside in G0, though in particular, yolk sac derived macrophages can proliferate ^1^. In addition tissue macrophages are also able to self-replenish, independent of circulating monoctyes ^2,3^. We previously observed cell cycle progression from G0 to G1 in monocyte-derived macrophages (MDM) stimulated by fetal calf serum (FCS) ^4^. This was mediated via the canonical mitogen-activated Ras/RAF/MEK/ERK signalling pathway, culminating in CDK4/6 activation and de-repression of Rb protein and transcription of cell cycle-associated proteins such as CDKs ^4,5^. The MEK/ERK pathway is evolutionarily conserved and involves feedback loops ^6^.

SAMHD1 is a deoxynucleotide-triphosphate hydrolase that lowers intracellular dNTP levels and works in opposition to ribonucleotide reductase (RNR) ^7–9^. This hydrolase activity is regulated by CDK1 phosphorylation of SAMHD1 at position T592, and during cell division CDK1 activity rises, thereby allowing accumulation of dNTP for S phase ^10^. We previously showed that macrophage cycle entry into G1 also correlates with an increase in CDK1 expression which phosphorylates SAMHD1 at position 592 and increases dNTP concentrations ^4^. SAMHD1 is also known as a restriction factor for virus replication of HIV-1 and DNA viruses, and thought to be mediated by decreasing the dNTP pool for viral DNA synthesis ^9,11,12^. Phosphorylation at T592 renders SAMHD1 inactive against HIV-1 ^10,13^. In support of an important role for SAMHD1 in virus restriction, the lentiviral accessory protein Vpx in SIV and HIV-2 have been shown to counteract and degrade SAMHD1 through proteasomal degradation ^14,15^. HIV-1 does not encode a Vpx like protein and bypasses SAMHD1 by infecting cells whilst they are in proliferative phases. Therefore the ability to infect non-dividing myeloid cells and T cells has been attributed to SAMHD1 ^14,16^. Importantly, infection of sentinel macrophages may contribute to viral persistence and pathogenesis ^17–19^.

We have subsequently sought to identify physiological factors that may impact macrophage cell cycle status; oxygen tension was of particular interest given the wide variability in oxygen tensions that macrophages encounter when passing through the circulation and across tissues ^20,21^. Indeed in areas where viruses such as HIV replicate, such as lymphoid tissue or the GI tract, tissue oxygen tension is 1-5% ^22,23^, in contrast to 11-14% in the circulation, and typical *in vitro* culture conditions at 21% oxygen. At low tissue oxygen levels, Hypoxia Inducible transcription Factors (HIFs) are stabilised, initiating a transcriptional programming response to adapt to changing oxygen levels ^24,25^. HIFs are ubiquitously expressed in all cells, but the main HIF isoforms, HIF1α and HIF2α, can be differentially expressed and can result in distinct transcriptional outcomes ^26^. However, both isoforms are regulated by PHDs (prolyl-hydroxylases 1-3) and FIH (factor inhibiting HIF), which are the oxygen sensors and act enzymatically in an oxygen-dependent manner to hydroxylate HIFα^27–29^. Prolyl hydroxylated HIFα acts as the recruitment signal for the Von Hippel-Lindau (VHL) E3 ligase, leading to HIFα proteasome-mediated degradation ^29^. However in low oxygen, PHD enzymatic activity is inhibited, leading to the stabilisation of HIFα followed by its translocation with HIF1β to the hypoxia response element (HRE) in the nucleus, cascading the activation of HIF-dependent gene expression ^30^. Asparagine hydroxylation of HIFα by FIH acts as negative feedback mechanism to control HIF-mediated transcription, and this results from the differential oxygen affinity between the PHDs and FIH ^31^.

Here we show that lowering oxygen tension to 1% (typical for tissues such as lymph nodes) results in a PHD-initiated cascade and a HIF2α-directed hypoxic transcriptional program in macrophages leading to reversible cell cycle progression. This response is accompanied by the activation of the canonical MEK-ERK pathway and expression of cell cycle-associated proteins including CDK1, which in turn deactivates the antiviral response of SAMHD1.

## Results

### Hypoxia drives reversible macrophage cell cycle progression

We isolated monocytes from blood of healthy individuals and differentiated them into monocyte-derived (MDM) macrophages using MCSF for 6 days (Figure 1A). The MDM were then placed in a hypoxia chamber at 1% oxygen. In parallel, identical wells of cells were placed in a standard tissue culture incubator supplied with 21% oxygen. We stained macrophages under 1% and 21% oxygen using a panel of myeloid antibodies that have been used to define M1 versus M2 macrophages (Supplementary Figure 1A). The expression of these markers did not indicate classical M1/M2 macrophage polarisation under hypoxic conditions. Phagocytosis is a core function of macrophages and we therefore assessed phagocytosis using the phrodo system ^32^. We observed an apparent heterogeneity in phagocytic activity and did not observe significant differences based on oxygen conditions across three donors (Supplementary Figure 1B).

**Figure 1:**
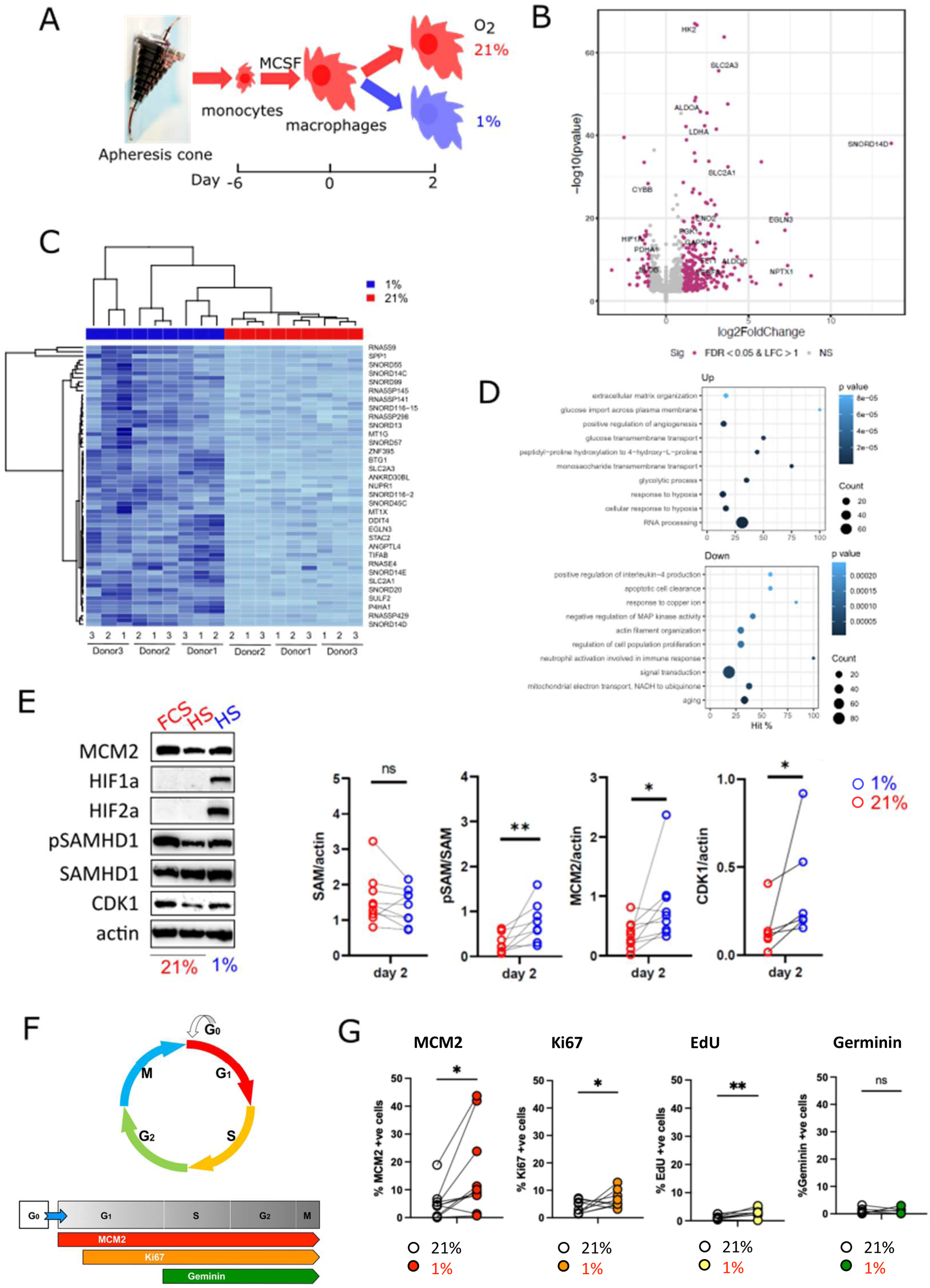
Terminally differentiated monocyte-derived macrophages (MDM) enter the cell cycle in hypoxia. A: a schematic diagram showing the experimental procedure. B: Volcano plot of differential mRNA expression in macrophages in hypoxia over normoxia. C: Heat map showing differentially expressed (DE) genes (FDR < 0.05, LFC > 2) between hypoxia and normoxia samples. Colour intensity is proportional to the Log2 Fold Changes. D: Functional-enrichment profiles in terms of Gene Ontology (GO)-based biological processes (BPs). E: Western blots of cell lysates from cells either stimulated with FCS or after two days of incubation in hypoxia were compared with those in normoxia. The antibodies used for blotting are listed by the left of the blots. FCS: fetal calf serum; HS: human serum. Quantitation of cell cycle markers (MCM2 and CDK1) and the phosphorylation of SAMHD1 (pSAM) and SAMHD1 were normalized with actin before plotting in pair. F&G: quantification of the cell cycle markers by immunofluorescence. Data represent at least 6 donors. Wilcoxon test used for statistical analyses with * p<0.5, ** p<0.1, *** p<0.01.

Bulk RNA seq analysis revealed that in these macrophages known HIF-driven genes were indeed transcriptionally regulated under low oxygen conditions (Figure 1B-D). Pathway analysis also showed enrichment of glycolytic genes and other canonical HIF target genes as expected (Figure 1B&C). Interestingly, genes involved in MEK/ERK pathway such as those encoding RAF, MEK, ERK, CDK4 and Cyclin D were expressed to higher levels in 1% oxygen (Supplementary Figure 2). In addition, the negative regulator to MAP kinase (ERK) (Figure 1D), which itself activates a plethora of downstream targets including the cell cycle regulating CDK1, was among the most downregulated genes. These observations, in conjunction with our previous study on FCS-stimulated MEK/ERK-induced cell cycle progression ^4^, prompted us to further examine the MEK/ERK pathway in the context of hypoxia. To further investigate whether hypoxia induces cell cycle progression, we lysed cells which had been incubated in either 1% or 21% oxygen for two days and performed western blot analyses to probe cell cycle-associated proteins. The cells treated at 1% oxygen displayed a robust expression of HIF1α and HIF2α, which were undetectable under the experimental setting in 21% oxygen tension (Figure 1E). We probed for MCM2 and CDK1 that we had shown previously to be reliable markers of cell cycle re-entry from G0 in FCS-stimulated MDM ^4^. Indeed, we observed an elevated expression of MCM2 as well as an up-regulation of CDK1 – the kinase responsible for SAMHD1 phosphorylation ^15^, similar to the observation for the MDM stimulated under FCS. We observed an elevated level of pSAMHD1 due to an elevated level of CDK1 expression while the overall expression of SAMHD1 was largely unaltered (Figure 1E).

Consistent with the western blot analysis, immunofluorescence staining on fixed cells post two day treatment revealed a marked increase in cell cycle-associated markers MCM2 as well as Ki67 and EdU (Figure 1F&G). Ki67 was detected at various stages of the cell cycle in cells exposed to 1% oxygen. EdU incorporation was marginally increased in 1% oxygen treatment, suggesting that synthesis is increased and replication stress was not induced. This is in contrast to more profound levels of hypoxia (<0.1%) where EdU incorporation is reduced, and replication stress is observed ^33^. The expression level of S/G2/M marker Geminin was not significantly altered, suggestive of cells not progressing into G2/M phase. Taken together we conclude that hypoxia drives MDM to re-enter cell cycle from G0 state, highly similar to our early studies using FCS as a stimulator for cell cycle progression in macrophages ^34^.

We next aimed to assess whether the cell cycle entry was reversible. We incubated differentiated MDM in 1% oxygen for two days before transferring them to an incubator supplying 21% oxygen and lysed the cells for different time intervals for western blotting (Supplementary Figure 3). We observed an acute and transient decrease in MCM2, CDK1 and pSAMHD1 after re-oxygenation along with a rapid loss of HIF1α and HIF2α. Interestingly at 30 mins post re-oxygenation we observed rebound expression of MCM2, CDK1 and pSAMHD1 but not HIFs. This rebound lasted at least 30 mins and had resolved by the final time point of 48 hours. We conclude that cell cycle changes in macrophages under hypoxic conditions are reversible and likely involve feedback circuits within the MEK/ERK axis that generates a temporary rebound in pERK leading to oscillatory pSAMHD1 expression after re-oxygenation.

### Macrophage cell cycle entry under low oxygen toggles SAMHD1 antiviral activity

We and others had previously observed that an increase in pSAMHD1 correlates with the alleviation of post-entry restriction of HIV-1 infection in myeloid cells ^4^. Notably this correlation does not hold for lentiviruses that encode Vpx protein, such as SIV or HIV-2, where SAMHD1 undergoes Vpx-mediated degradation via proteasomes ^14^. As pSAMHD1 levels were increased in 1% oxygen-treated MDMs, we hypothesised that the restriction of HIV-1 infection in these macrophages should be alleviated and hence an increase in viral replication, which should contrast with the level of infection in SIV or HIV-2. To test this hypothesis, we produced HIV-1, SIV or HIV-2 pseudotyped with VSV-G glycoproteins together with packaged genomes that once integrated into the target cells express GFP for ease of quantitation of infected cells. Western blot analysis showed that the 1% oxygen resulted in upregulation of CDK1 which drives an enhanced pSAMHD1 expression (Figure 2B) and this increase correlates with up to a 5-fold increase in HIV-1 infection (Figure 2A). As predicted, this increase in infectivity is not observed for SIVmac or HIV-2, where SAMHD1 is degraded (Figure 2B & Supplementary Figure 4). We further knocked down SAMHD1 and found the infectivity was increased in 1% oxygen condition (Figure 2C). Thus the toggling of SAMHD1 antiviral activity serves as a surrogate marker for macrophage cell cycle reentry, thereby also validating our observations for cell cycle markers (MCM2, Geminin, EdU) and associated proteins/transcripts.

**Figure 2:**
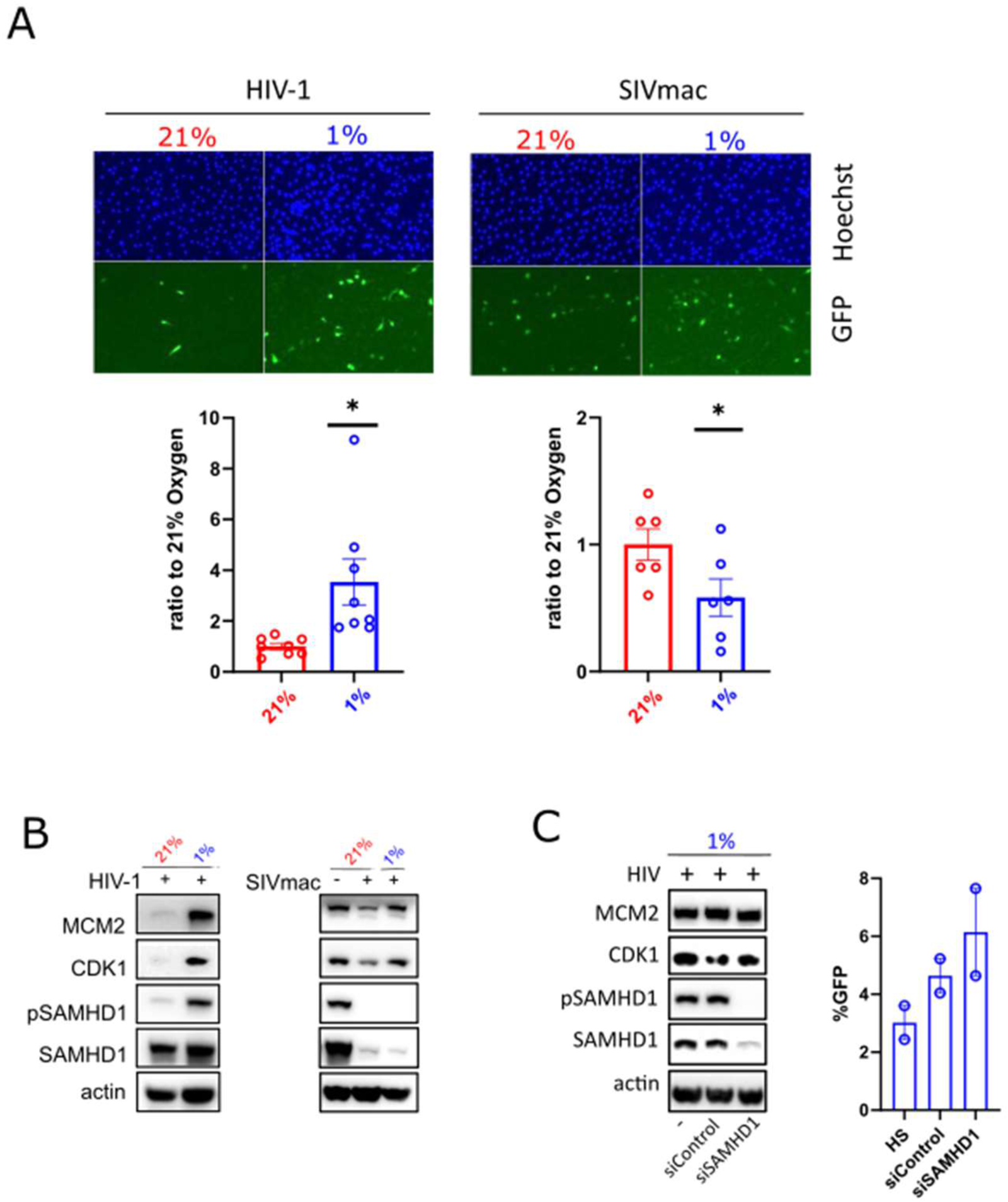
Hypoxia increases susceptibility to virus infection due to cell cycle progression and SAMHD1 deactivation. A: fluorescent images of HIV-1 and SIVmac infection with Hoechst for nuclei and GFP for virus infection in both normoxia and hypoxia are shown. The number of GFP-positive cells were quantitated and normalized to the total number of cells to give a single-round infection rate. The rates were then normalized with those in normoxia conditions for either HIV-1 or SIVmac infection. Duplicate infections are plotted for each donor with at least three donors in each condition. One sample t test is used for statistical analyses with * p<0.5. B: Western blots for conditions in A are shown. C: SAMHD1 KD and infectivity. The antibodies used for blotting are shown by the left of the blots.

### MEK/ERK inhibitors abrogate cell cycle changes induced by low oxygen conditions

The MEK/ERK pathway is enriched in our transcriptomics study and has been shown to be involved in the FCS-induced cell cycle progression ^4^. Hence we hypothesised that MEK/ERK pathway might be implicated in hypoxia-driven cell cycle progression. We took advantage of clinically proven antagonists against different targets in this pathway to examine the specificity of this pathway in driving the cell cycle progression in macrophages (Figure 3A). We found the inhibitors indeed efficiently and specifically blocked the activation cascade in MDM as the inhibitors to the upstream RAF (Tak632 and AZ628) completely abolished the phosphorylation on ERK therefore the abrogation of expression in CDK1 (Figure 3B). MEK inhibitor U0126 also manifested inhibition in phosphorylation of ERK but not the CDK4/6 inhibitor palbociclib which acts downstream of ERK but upstream of CDK1 expression. Nevertheless, all the MEK/ERK inhibitors led to a drastic decrease in CDK1 expression. Since signal transduction is converged at ERK, we chose to focus on using U0126 and palbociclib, one upstream and one downstream of ERK for further studies of the regulation of MEK/ERK pathway in hypoxia. MDMs were incubated in 1% oxygen for 12 hrs prior to the addition of U0126 or palbociclib for 2 hrs; this was followed by transduction using pseudotyped viruses bearing VSV-G for single-round infections. The treatment with U0126 completely abrogated the phosphorylation of ERK which then culminated in decreased CDK1 expression in 1% oxygen, resulting in a diminished level of pSAMHD1 (Figure 3C). However, the treatment of palbociclib maintained the expression level of pERK but led to a decrease in CDK1 and thus phenocopied the effect seen in the U0126 treated cells. In accordance with the observed expression profile for pSAMHD1 by western blotting, the infectivity of HIV-1 was completely abolished by either U0126 or palbociclib, confirming the role of hypoxia in regulating the MEK/ERK pathway, cell cycle re-entry and HIV-1 restriction in macrophages (Figure 3C).

**Figure 3:**
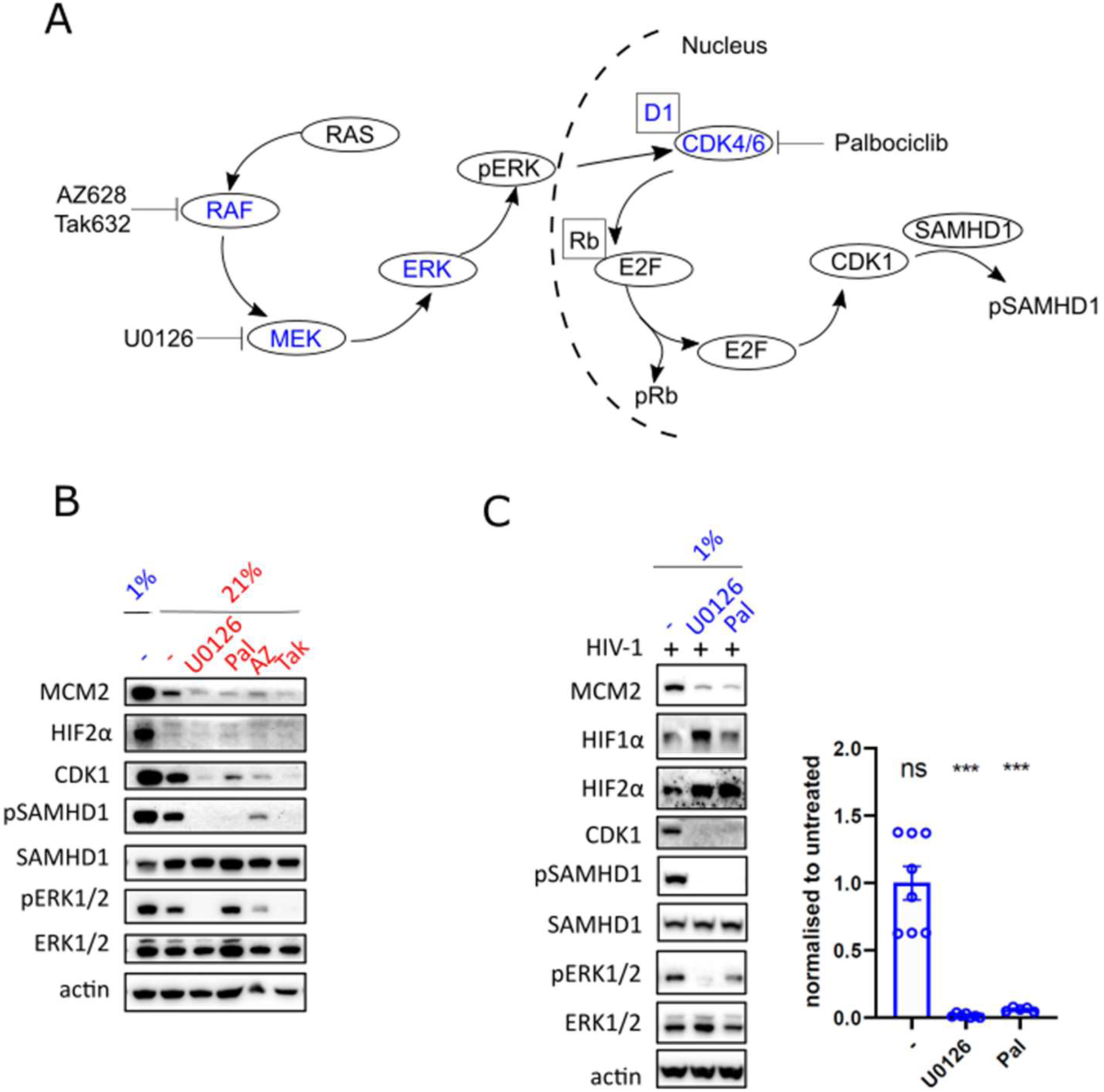
MEK/ERK pathway is involved in hypoxia-induced cell cycle progression. A: A diagram showing the MEK/ERK pathway and the specific inhibitors to different targets within this pathway. Targets depicted in this diagram with increased levels of mRNA expression (supplementary figure 2) in hypoxia are highlighted. B: verification of the drug specificity in normoxic condition. Inhibitors used were U0126 (10 uM), Palbociclib (10 uM), Tak (2 uM) and AZ (1 uM). Representative western blot of cell lysates which was harvested post two days incubation in hypoxia or with an addition of the inhibitors in normoxia. The antibodies used for blotting are shown by the left of the blots. C: Representative western blot of lysates from the cells treated with U0126 and Pal or without followed by HIV infection for two days in hypoxia. The number of GFP-positive cells was quantitated and normalized to the total number of cells to give a single round infection rate. The rates were then normalized with those untreated for HIV-1 infection. Duplicate infections are plotted for each donor with at least three donors in each condition. One sample t test is used for statistical analyses with *** p<0.01.

### Mouse peritoneal macrophages enter the cell cycle in low oxygen conditions

We previously observed mouse peritoneal macrophages stimulated by FCS could re-enter cell cycle ^4^. To examine whether the low oxygen tension-driven cell cycle re-entry is specifically for human cells, we sought to examine this by using mouse peritoneal macrophages exposed to 1% oxygen. As a control to the enhanced HIV-1 infectivity in 1% oxygen, we additionally collected peritoneal macrophages from SAMHD1 KO mice which should de-suppress the restriction to infection. The cells from the same WT or SAMHD1 KO animal were split into two aliquots for culture in 21% or 1% oxygen. Western blot analysis of the lysates showed that, although there was variability between cells from the WT animals, KO cells displayed a significant increase in MCM2 expression in 1% oxygen (Figure 4A). Consistent with the observation that pERK is increased in this hypoxic condition in human cells, we found that peritoneal macrophages from both WT and KO animals had elevated levels of pERK in 1% oxygen, demonstrating that hypoxic regulation of the MEK/ERK pathway is evolutionarily conserved. As expected, SAMHD1 was absent from the KO cells and infection using VSV-G pseudotyped HIV virus showed a 20-fold increase in infection in SAMHD1 KO cells compared to WT cells (Figure 4B). The infection in WT was negligible (0.1%) due to the presence of active SAMHD1. However, when the cells were exposed to 1% oxygen a small but statistically significant 3-fold increase in infection was noted. This is in agreement with our data from human MDMs and shows that hypoxia drove macrophages into a non-G0 stage which correlated with an increase in susceptibility to pseudotyped HIV-1.

**Figure 4:**
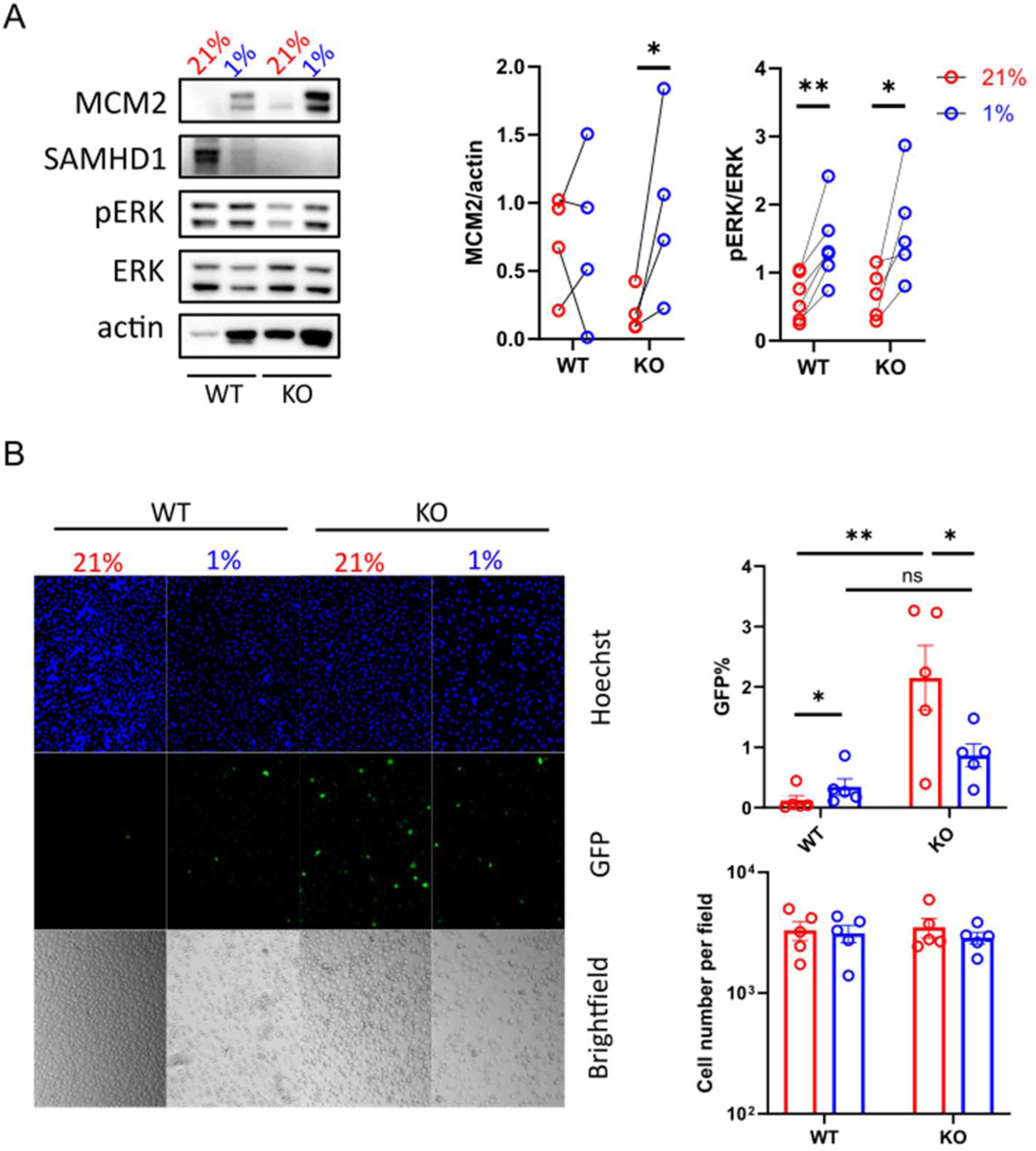
Hypoxia induced cell cycle progression in mouse peritoneal macrophages. A: Representative western blot of WT and SAMHD1 KO cell lysates harvested post two days in hypoxia or normoxia. The antibodies used for blotting are shown by the left of the blots. Quantitation of MCM2 and the ratio of pERK/ERK are shown on the right. B: WT and SAMHD1 KO perineal macrophages were infected with HIV either in hypoxia and normoxia. The number of GFP positive cells was quantitated and normalized to the total number of cells to give a single round infection rate. The rates were then normalized with those untreated for HIV-1 infection.

### PHD inhibitors mimic the effect of low oxygen on macrophage cell cycle progression

Having demonstrated that both MDM and peritoneal macrophages from mice can be driven to enter the cell cycle in hypoxia, we next sought to determine whether manipulating the HIF pathway using pharmacological inhibitors in 21% oxygen was sufficient to replicate our findings in 1% oxygen (Figure 5). As there are several classes of clinically proven or under clinical investigation inhibitors targeting different stages of HIF-mediated pathway, we selected well-documented PHD inhibitors. This class of drugs is used in treatment of anemia associated with renal anemia ^35^. We observed that drugs targeting PHD activity (daprodastat) increased the stabilisation of HIF (Figure 5A). Interestingly daprodustat-treated cells manifested a significant increase in MCM2 expression, suggesting the presence of those drugs drives PHD-dependent cell cycle entry through stabilization of HIF1α or HIF2α. Strikingly, the treatment of daprodustat in normal oxygen levels led to an elevated level of pSAMHD1 which correlated with ∼5 fold increased susceptibility to HIV-1 infection. This increased infectivity was abrogated in the additional presence of either U0126 or palbociclib (Figure 5B&C), recapitulating the phenotype observed in hypoxia (Figure 2).

**Figure 5:**
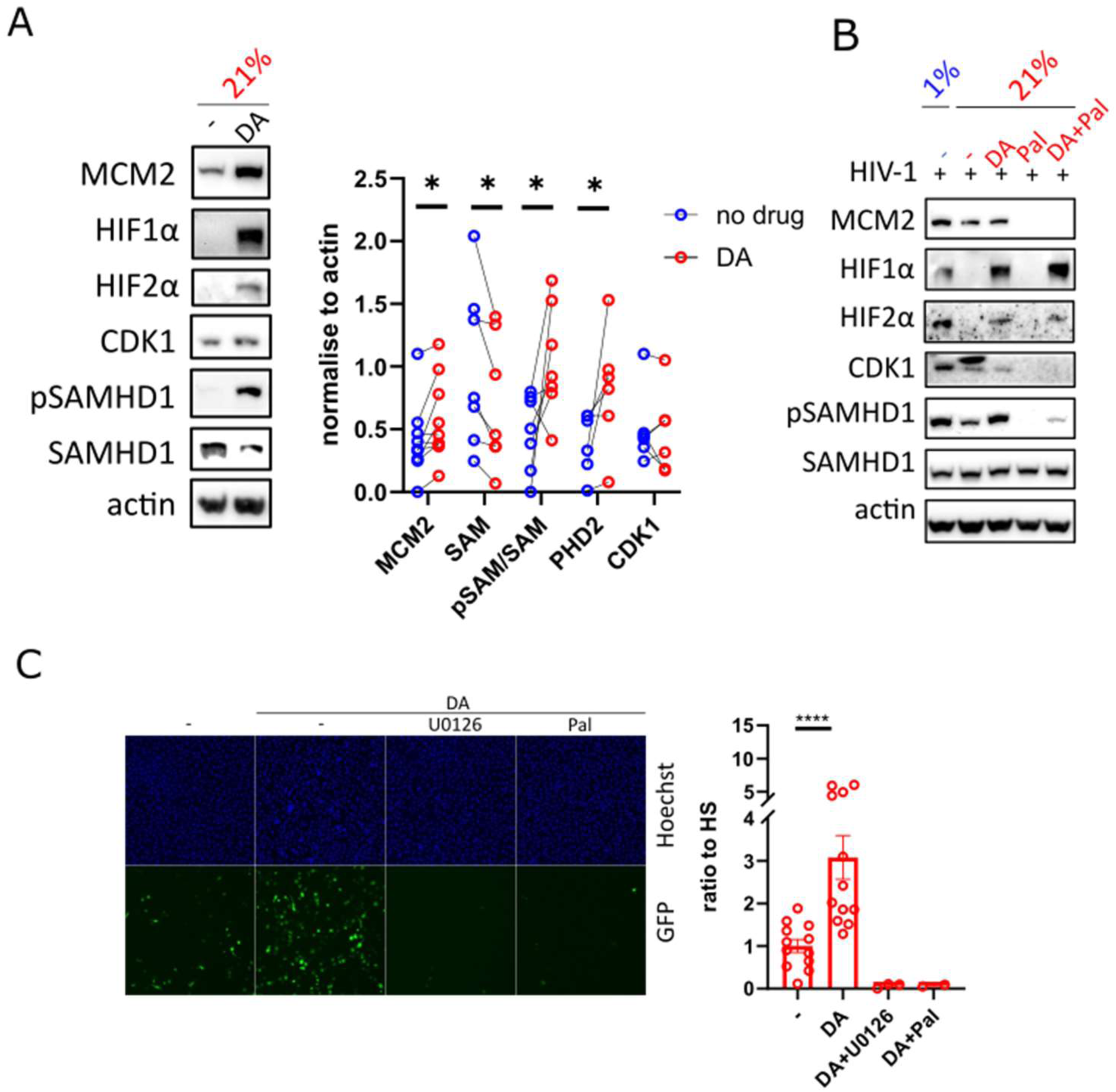
PHD inhibition mimics hypoxia and drives macrophage cell cycle progression. A: Representative western blot of cell lysates harvested post two days of treatment of Daprodastat (DA) in normoxia. The antibodies used for blotting are shown by the left of the blots. The quantitation of the protein bands was densitometrically quantified and plotted between drug-treated and untreated samples. Each pair represents one prep from one donor. B: Representative western blot of cell lysates with MEK/ERK inhibitors followed by the HIV infection. C: fluorescent images of HIV-1 infection with Hoechst for nuclei and GFP for virus infection in both drug-treated and untreated cells are shown. The number of GFP positive cells was quantitated and normalized to the total number of cells to give a single-round infection rate. The rates were then normalized with those in the untreated condition. Duplicate infections are plotted for each donor with at least three donors in each condition. One sample t test is used for statistical analyses with * p<0.5. **** p <0.001

### Macrophage cell cycle progression is dependent on HIF2**α** activity

We next sought to investigate whether the cell cycle re-entry is dependent on HIF transcription factors. We decided to examine the involvement of HIF2α using the clinically approved selective HIF-2α inhibitor, PT-2385. This inhibitor disrupts the interaction between HIF2α and HIF1β leading to reduced expression of HIF2α-dependent HIF target genes ^36^. Indeed, the presence of PT-2385 did not affect the expression of unrelated genes (BAP1) or genes mainly regulated by HIF1α (PDK-1 and PGK-1) while PAI-1, a gene mainly targeted by HIF2α, was specifically reduced in a dose-dependent manner (Figure 6A). We also observed a dose-dependent decrease in MCM2 expression in the presence of PT-2385 under hypoxia (Figure 6B), indicating that PT-2385 effectively blocks the cells from re-entering the cell cycle in hypoxia. Accordingly, CDK1 expression was decreased with PT-2385 treatment accompanied by a reduced level of pSAMHD1. Since pERK functions upstream of CDK1 expression (Figure 3A), we reasoned that the level of pERK may be affected in the presence of PT-2385. Indeed while total ERK was unaltered in the presence of the drug (Figure 6C), a statistically significant two-fold reduction in pERK expression was evident (Figure 6D). Additionally, PT-2385 significantly decreased the susceptibility to HIV-1 infection (Figure 6E), whilst cell numbers were largely maintained (Figure 6F).

**Figure 6:**
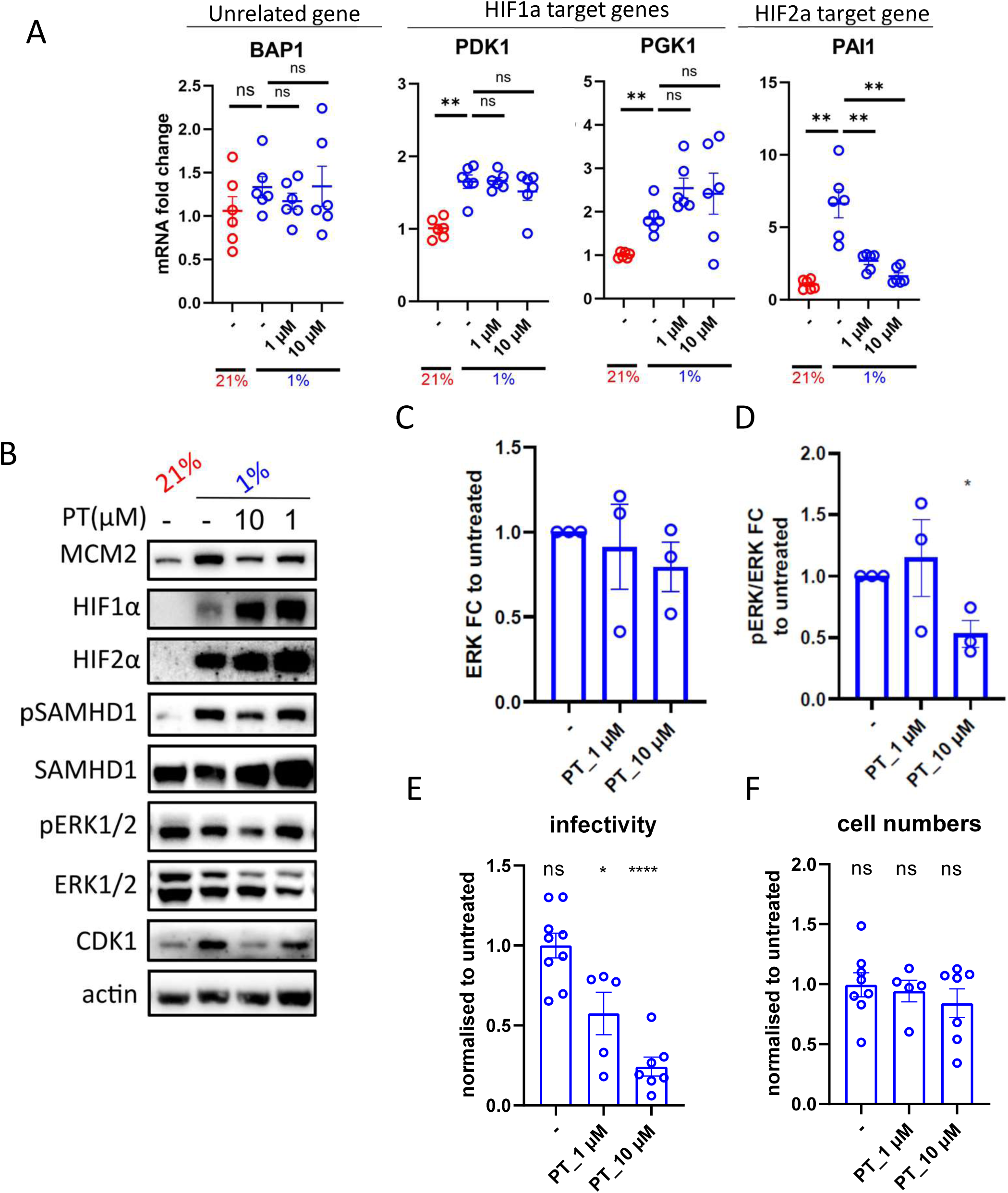
HIF2a is required for hypoxia-induced cell cycle progression. A: MDM were harvested two days post addition of PT-2385 HIF2a antagonist (PT) in hypoxia before harvested for extraction of RNA followed by RT-PCR analyses. B: western blot of cell lysates harvested post two days addition of PT in hypoxia in comparison to the untreated samples either in hypoxia or normoxia. The antibodies used for blotting are shown to the left of the blots. C&D: pERK and ERK bands from the drug-treated and untreated samples were quantified by densitometry, normalized against actin and plotted. E&F: The number of GFP-positive cells was quantitated and normalized to the total number of cells to give a single-round infection rate. The rates were then normalized with those untreated. Duplicate infections are plotted for each donor with at least three donors in each condition. One sample t-test is used for statistical analyses with * p<0.5.

To further investigate whether HIF2α was also involved in the cell cycle progression induced by PHD inhibitors, we sequentially treated the cells with daprodustat followed by PT-2385. As predicted, daprodustat treatment induced cell cycle re-entry that was abrogated with PT-2385 (Supplementary Figure 5). Given both CDK1 and pSAMHD1 were reduced we hypothesised that the level of pERK expression was lowered. Western blot analyses on multiple donors indeed showed that the total ERK was unaltered (Supplementary Figure 5B) but that the ratio of pERK/ERK was reduced in the presence of the higher dose of HIF2α antagonist (Supplementary Figure 6C), which correlated with decreased HIV-1 infectivity (Supplementary Figure 6D&E). Taken together our data suggest that HIF2α is necessary and sufficient to account for the observed hypoxia-driven cell cycle progression as well as the PHD inhibitor-induced cell cycle response in macrophages.

### Tumour-associated macrophages exhibit hypoxia and cell cycle progression signatures

Hypoxia is one of the hallmarks of tumour growth environment where tumour-infiltrating macrophages are also found. Lung adenocarcinomas are known to have mutations in the MEK/ERK pathway. Given the interplay between the MEK/ERK pathway, hypoxia and cell cycle re-entry in MDM, we sought to explore the presence of hypoxia and cell cycle-related gene signatures within macrophages *in vivo* from two independent single-cell RNA sequencing data of lung adenocarcinoma^58,59^. For both datasets, we identified different clusters of tissue-resident and monocyte-derived macrophages (MDMs), monocytes, dendritic cells, and alveolar macrophages (Fig. 7A, Supplementary Fig. 6A) based on cell type marker genes (Fig. 7B, Supplementary Table 1).

**Figure 7:**
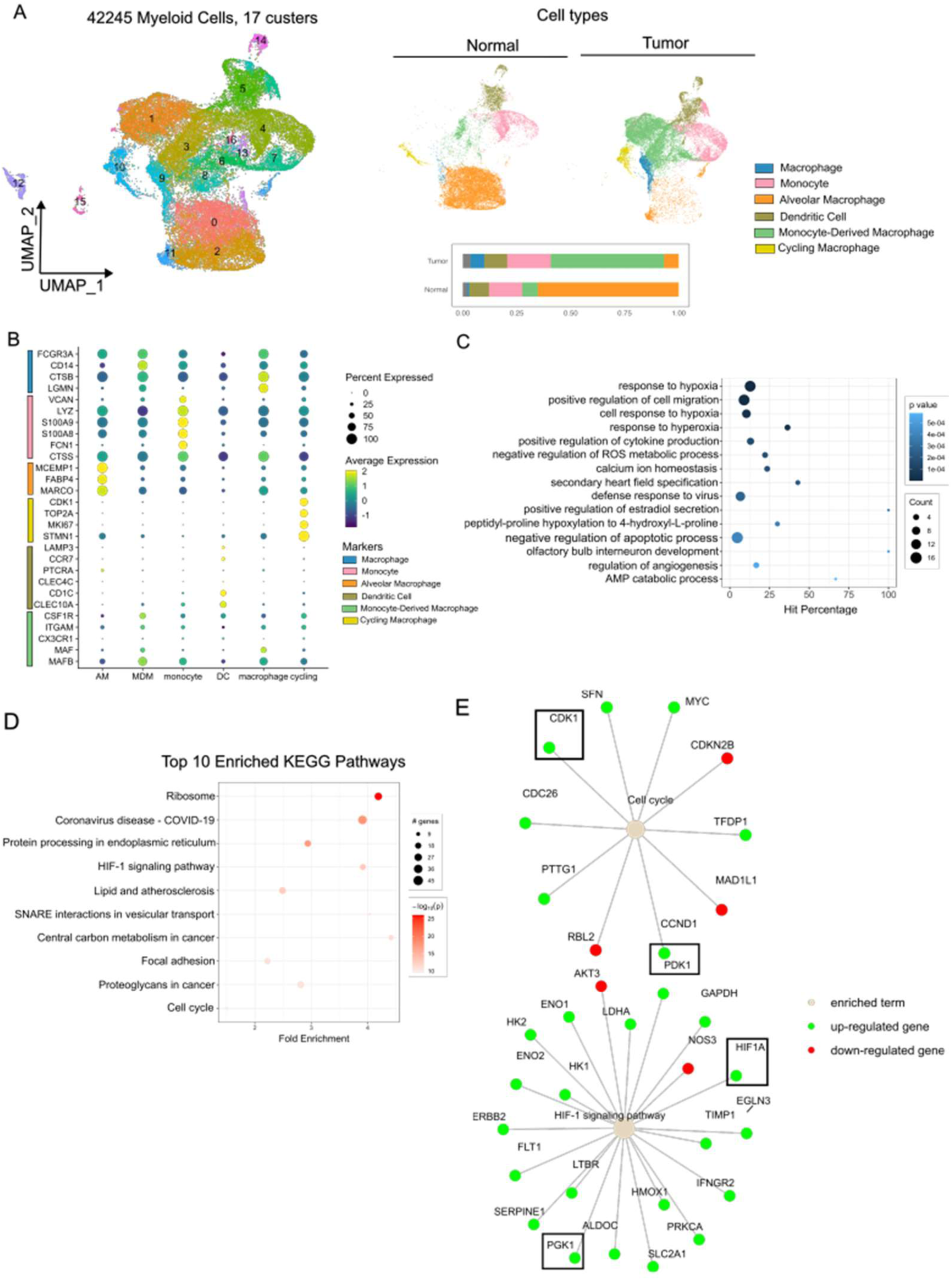
Tumour-associated macrophages exhibit hypoxia and cell cycle progression signatures. Using the scRNA seq dataset from^58^. A. UMAPs based on the top 15 principal components of all myeloid single-cell transcriptomes split by tissue type, color-coded by cell cluster. B. Average gene expression of selected marker genes for myeloid cell clusters. C. Functional-enrichment profiles in terms of Gene Ontology (GO)-based biological processes (BPs). The hit percentage is the ratio of the DEG number and the number of all genes in a GO term. D. KEGG pathway enrichment analysis of DEGs in tumor MDMs. The enrichment factor is the ratio of the DEG number and the number of all genes in a certain enrichment pathway. The dot size denotes the number of DEGs, while colours correspond to the log of the adjusted p-value range. E. Term-gene graph of cell-cycle and hypoxia related KEGG pathways.

We noted quantitative shifts in the cellular composition of the tumour immune microenvironment (Fig. 7A). Among myeloid cell types, MDMs and dendritic cells were increased, while tissue-resident macrophages and monocytes were decreased, which we also observed in an independent second dataset (Supplementary Fig. 6A).

We then conducted pseudobulk differential expression analysis on the tumour MDM cluster. KEGG Pathway analysis revealed enrichment of Cell Cycle and HIF1-signalling pathway (Fig. 7C). Gene Ontology (GO) analysis showed enrichment of hypoxia-related GO terms (Fig. 7D). Genes involved in cell cycle such as CDK1, CDC26, and MYC are enriched (Fig. 7E). Consistent with gene expression analysis from MDM in 1% oxygen (Fig.6), HIF1α-regulated genes PDK1 and PGK1 are upregulated in tumour MDMs (Fig. 7E, black box). We further probed for MCM2 and CDK1 expression, which are reliable markers of cell cycle re-entry of MDM from G0. Indeed, we observed an upregulation of MCM2 and CDK1 in TAMs (Fig. 7 & Supplementary Fig. 6E, black boxes). In the second independent dataset, we also observed enrichment of cell cycle pathways (Supplementary Fig.6C) and hypoxia-related GO terms (Supplementary Fig.6D, E). In addition, we observed upregulation of genes involved in MEK/ERK pathway such as MAPK12 and MYC (Supplementary Fig.6E, black boxes). Taken together we show enrichment of hypoxia and cell cycle related gene signatures in tumour associated MDMs at single cell level.

## Discussion

Cell cycle regulation not only impacts the process of cell division but also inflammatory responses ^37,38^. High dNTP levels are detrimental for cells, both due to increased likelihood of parasitisation by DNA viruses and other micro-organisms, and possibly higher rates of spontaneous mutation following DNA damage and repair. Although macrophages were traditionally viewed as non-proliferating cell types, more recent insights have shown that yolk sac derived macrophages in tissues such as brain and gut have significant proliferative potential ^1^, and monocyte derived tissue macrophages are able to proliferate in response to specific signals ^39^. These insights have significant implications for health and disease ^40^. While our previous studies have demonstrated that MDM can be stimulated by FCS and re-enter the cell cycle ^4^, we were interested in how monocyte-derived macrophages respond to low oxygen tension, which reflects physiological conditions where those cells reside *in vivo*. Here we report that hypoxia can drive cell cycle entry in monocyte-derived macrophages. This was somewhat surprising as hypoxia was previously shown to arrest the cell cycle in numerous other cell types ^41,42^. In contrast, HIF2α is the key driver in clear cell renal cell carcinoma where VHL is dysregulated ^43^ and there is an emerging recognition that HIF1 and 2 can impact cell cycle differentially in the context of neoplasia ^44^. Therefore it is intriguing and important that the same physiological stimulus can drive opposing responses based on cell type, and these may be driven through direct and indirect interactions with oncoproteins and tumour suppressors such as Myc and p53 respectively ^44^.

Cell cycle progression resulted in modestly increased incorporation of EdU. Surface expression of classical macrophage markers such as CD86 did not change significantly in response to hypoxia (Supplementary Figure 1A), and phagocytic activity was not substantially altered in hypoxic macrophages (Supplementary Figure 1B). We have demonstrated from three lines of evidence that a subpopulation of macrophages re-enter into the non-G0 stage by a hypoxia-triggered canonical mitogen-activated proliferation cascade in macrophages, involving MEK-ERK and downstream CDK4/6-Cyclin D, with phosphorylation of SAMHD1. Firstly, our transcriptomic analysis highlighted, and transcript analysis by qRT-PCR confirmed, genes involved in the MAPK pathway are highly differentially expressed in hypoxia. Negative regulators of ERK were also downregulated additionally implicating this pathway in the response to low oxygen tension. Secondly, pharmacological inhibition on the intermediate targets involved in the MEK/ERK pathway completely abolished the low oxygen tension-induced cell cycle re-entry phenotype. Thirdly, the addition of an antagonist to HIF2α either in hypoxia or PHD inhibitor-driven HIF stablisation in normoxia prevents cells from re-entering from G0, with a decrease in ERK phosphorylation and CDK1 expression.

We found that the cell cycle changes were reversible within minutes following restoration of oxygen tension. Interestingly at 30 mins post re-oxygenation we observed rebound expression of MCM2, CDK1 and pSAMHD1, but not HIFs (Supplementary Figure 3). This rebound lasted at least 30 minutes and had resolved by the final time point of 48 hours. We conclude that cell cycle changes in macrophages under hypoxic conditions are reversible and likely involve feedback circuits within the MEK/ERK axis that generate an oscillatory rebound ‘hypoxia-like’ response after re-oxygenation that is HIF independent. This may be similar to work showing paradoxical ERK activation in the presence of RAF inhibitors ^45^. Interestingly we observed that ERK1/2 is upregulated in hypoxia which can be abrogated in the presence of a HIF antagonist – suggesting ERK may be transcriptionally regulated by HIF2a, consistent with the identification of an HRE upstream of the ERK promoter ^46^.

Mechanistically, we identified HIF2α as the key mediator. This was supported by the use of a specific inhibitor of HIF2α (PT-2385) that was able to completely abrogate the effect of hypoxia-driven cell cycle re-entry. PT-2385 blocked the upregulation of HIF2α specific transcripts and partially blocked ERK phosphorylation, suggesting that the pathway connecting hypoxia to cell cycle is proximal to MEK, with possible involvement of factors upstream of MEK such as RAF, RAS and EGFR. PHD3 is known to be highly upregulated in hypoxia and is known to negatively regulate the level of EGFR at the cell surface in certain conditions ^47,48^. However, we were unable to assess the potential role of EGFR in macrophages due to the low abundance of the protein as estimated by western blot (data not shown).

The dNTP hydrolase and antiviral restriction factor SAMHD1 is known to be regulated by cell cycle ^4,49^. The block to HIV-1 infection is governed by CDK1-dependent phosphorylation at T592 ^10,13^. SAMHD1 is phosphorylated when macrophages re-enter the cell cycle from G0 ^4^. Although there are instances where cell cycle-associated proteins such as cyclin D3 can restrict viruses independent of cell cycle status ^50^, SAMHD1 restriction of HIV-1 is closely related to cell cycle status. As expected, the primate lentivirus SIVmac, encoding an accessory protein Vpx that degrades SAMHD1, infected macrophages to a similar extent under hypoxic and normoxic conditions, suggesting SIVmac is likely to be insensitive to macrophage cell cycle status. These data reinforced that cell cycle status has significant impacts on susceptibility to viruses that do not have a means of antagonising the activity of SAMHD1, for example HIV-1. Here we observed that the cell cycle entry associated with hypoxia, or PHD inhibition, results in phosphorylation and deactivation of SAMHD1 antiviral activity in macrophages. This phenotype could be blocked by pharmacological MEK inhibition as well as CDK4/6 inhibition (Figure 3B&C), indicating that the proliferation pathway activated by hypoxia was similar to that previously elaborated in macrophages ^4^. As predicted, HIV-2 – a lentivirus that encodes the SAMHD1 antagonist Vpx – is not impacted by hypoxia-induced cell cycle entry (Supplementary Figure 4), confirming the primacy of SAMHD1 in the restriction of HIV-1 infection in macrophages. These data have implications for the maintenance and expansion of HIV-1 reservoirs in lymphoid tissue where oxygen tensions are estimated to be between 1-5%. Drugs that selectively block the link between HIF2 and cell cycle progression could be used therapeutically.

Using single cell transcriptomic analysis of two independent cancer datasets, we observed an association between hypoxia and cell cycle progression transcriptional signatures in tumour associated macrophages (TAM), consistent with our model of hypoxia driving cell cycle progression. Moreover, HIF2α has been reported to be particularly highly expressed in tumour-associated macrophages (TAM) ^51^ .TAM-associated HIF2α has also been correlated with angiogenesis in breast cancer ^52^. This may be related to the fact that anaemia during cancers is common ^53^and reduced oxygen delivery to tissues during anaemia ^54^ could result in HIF2α activation, and macrophage cell cycle entry.

Inhibition of cell cycle is a central mechanism of cancer therapies, and future work should include an assessment of the relationship between TAM cell cycle inhibition and their modulation of the micro-environment, including anti-tumour activity and risk of metastasis. As deactivation of SAMHD1 in hypoxic macrophages renders them hyper-susceptible to lentiviral transduction, there is potential for efficient lentiviral gene delivery to target TAM and reverse the immune suppressive / metastatic environment.

## Limitations

Our study was limited by use of *ex vivo* cells and lack of *in vivo* functional data. We did however demonstrate the relationship between hypoxia, HIF2α and cell cycle re-entry in both tissue macrophages from mice as well as human monocyte-derived macrophages. In summary, we show that low oxygen tension results in a PHD-initiated cascade and a HIF2α-directed hypoxic transcriptional program in macrophages and reversible cell cycle re-entry. This response is the result of the activation of the canonical MEK-ERK pathway. Our work paves the way for further exploration of the impacts of oxygen tension on macrophage functions in inflammation, immunity, neoplasia and viral reservoirs.

## Acknowledgements

This work was funded by the Lister Institute and a Wellcome Senior Fellowship to RKG (WT108082AIA), as well as the UK Medical Research Council (MRC core funding of the MRC Human Immunology Unit; J.R.). This research was supported by the NIHR Cambridge Biomedical Research Centre (NIHR203312). The views expressed are those of the authors and not necessarily those of the NIHR or the Department of Health and Social Care’. We would like to thank Prasanti Kotagiri, Kenneth GS Smith and Paul Lyons.

## Figure legends

**Supplementary Figure 1: Macrophage surface markers and phagocytosic activity under 1% oxgen tension.** A: MDMs treated in 21% and 1% oxgen tension were stained with relative antibodies against surface markers or arginasel before analysis by FACS. B: Examples of Immunofluorescence images of phagocytosis analysis by pHrodo. The number of pHrodo-positive cells between 1% and 20% oxygen level was compared and plotted.

**Supplementary Figure 2: Transcriptomic signatures associated with hypoxia in macrophages** A: PCA bi-plot showing clustering of the different transcriptional profiles in 1% vs 21% oxygen exposed samples. B: KEGG pathway enrichment analysis of DEGs in macrophages exposed to 1% oxygen level. The enrichment factor is the ratio of the DEG number and the number of all genes in a certain enrichment pathway. The dot size denotes the number of DEGs, while colours correspond to the log of the adjusted p-value range. C: qRT-PCR verification of the target genes involved in HIF and MEK/ERK pathways.

**Supplementary Figure 3: The 1% oxygen-induced cell cycle progression in macrophages is reversible.** MDM had been incubated in a 1% oxygen chamber for two days before being transferred to an incubator supplying 21% oxygen. Western blots of cell lysates collected at different time intervals were compared with those in non-drug treated either 21% or 1% oxygen treated samples, or with U0126 in 1% oxygen condition. The antibodies used for blotting are listed to the left of the blots.

**Supplementary Figure 4: Single-round infectivity in macrophages is unaltered for HIV-2.** Immunofluorescence analysis was used to quantify the number of GFP-positive cells and normalized to the total number of cells to give a single-round infection rate. The rates were then normalized with those in 21% oxygen level for HIV-2 infection.

**Supplementary Figure 5: HIF2α is required for DA-induced cell cycle progression in 21% oxygen tension.** A: MDM were treated with DA before the addition of PT-2385 HIF2α antagonist (PT) in 21% oxygen level. Cells were lysed post two days of treatment for western blot analyses. The antibodies used for blotting are shown by the left of the blots. C&D: pERK and ERK bands from the drug-treated and untreated samples were densitometrically quantified, normalized against actin and plotted. E&F: The GFP-positive cells were quantitated and normalized to the total number of cells to give a single-round infection rate. The rates were then normalized with those untreated. Duplicate infections are plotted for each donor with at least three donors in each condition. One sample t-test was used for statistical analyses with * p<0.5.

**Supplementary Figure 6: Tumour-associated macrophages exhibit hypoxia and cell cycle progression signatures. Analysis of scRNA seq data from^59^:** A. UMAPs based on the top 15 principal components of all myeloid single-cell transcriptomes split by tissue type, color-coded by cell cluster. B. Average gene expression of selected marker genes for myeloid cell clusters. C. Functional-enrichment profiles in terms of Gene Ontology (GO)-based biological processes (BPs). The hit percentage is the ratio of the DEG number and the number of all genes in a GO term. D. KEGG pathway enrichment analysis of DEGs in tumor MDMs. The dot size denotes the number of DEGs, while colours correspond to the log of the adjusted p-value range. E. Term-gene graph of cell-cycle and MAPK signalling pathways.

## Materials and Methods

### Cells, plasmids and viruses

293T cells were cultured in DMEM supplemented with 10% FCS. MDM were cultured in RPMI 1640 medium supplemented with 10% human serum (Sigma). VSVDG pseudotyped HIVD1 GFP virus was produced by transfecting of 293T with GFPDencoding genome CSGW, packaging plasmid p8.91 and pMDG as previously described ^56^. VSVDG HIVD2 GFP virus was produced as previously described ^57^, and VSVDG, SIVsmE543 GFP virus was a kind gift from G. Towers (UCL).

### Inhibitors and antibodies

Kinase inhibitors used are CDK4/6 inhibitor (PD 0332991, Palbociclib, Sigma); MEK/ERK inhibitor U0126 (Calbiochem); RAF (TAKD632), AZ628; Daprodastat; PT-2385; Antibodies used were as follows: antiDSAMHD1 (ab67820, Abcam, UK), anti-betaDactin (ab6276, abcam, UK); mouse antiDMCM2 (BMD28, BD Biosciences, UK); pSAMHD1; CDK1; ERK1/2; pERK1/2; anti-HIF1α; anti-HIF2α; antiDGeminin (NCLDLDGeminin, Leica).

### Monocytes isolation and differentiation

PBMC were prepared from apheresis cones from NHS Blood Center Cambridge by densityDgradient centrifugation (Lymphoprep). MDM were prepared by adherence with washing of nonDadherent cells after 2 h, with subsequent maintenance of adherent cells in RPMI 1640 medium supplemented with 10% human serum (Sigma) and MCSF (10 ng/ml) for 3 days and then differentiated for a further 3 days in RPMI 1640 medium supplemented with 10% human without MDCSF. The cells are then incubated in either 21% or 1% oxygen tension for the remaining period of experiments.

### VSV-G pseudotyped virus single-round infection

VSVDGDpseudotyped HIVD1, HIV-2 or SIVmac GFP expressing viruses were added to MDM or tissueDresident macrophages from mice, which had been exposed to relevant oxygen tension incubator overnight. 2 h post-incubation, the inoculum was removed and cells were washed once in culture medium. This was left for 36 hr post infection before the cells were stained by Hoechst for nuclei. The percentage of infected GFP-expressing cells versus total cells was determined 48 h postDinfection by EVOS and analysed by using ImageJ software.

### Mice

Mice were housed and bred under standard conditions at the University of Oxford Biomedical Services Animal Facilities. *Samhd1^-/-^* mice were described before ^55^. All mice were on the C57BL/6 background and were 6-7 weeks old; male and female animals were used. This work was performed in accordance with the UK Animal (Scientific Procedures) Act 1986 and institutional guidelines for animal care. This work was approved by project licenses granted by the UK Home Office (PPL number PP5102963) and was also approved by the Institutional Animal Ethics Committee Review Board at the University of Oxford. Peritoneal macrophages were obtained after animals were humanely killed by washing the peritoneal cavity with 5 ml RPMI containing 10% FCS.

### SAMHD1 KD in MDM

This was done as previously described ^34^. Briefly, 1 × 10^5^ MDM differentiated in MCSF for 4 days were transfected with 20 pmol of siRNA (LD013950D01, Dharmacon) using Lipofectamine RNAiMAX Transfection Reagent (Thermofisher). Transfection medium was replaced after 18 h with RPMI 1640 medium supplemented with 10% human serum and cells cultured for additional 3 days before pseudotyped virus infection.

### Bulk RNAseq from MDM

Bulk RNA sequencing of cells was performed as described in (Kotagiri, Mescia et al. 2022) with adaptions. In brief, RNA was purified using the RNeasy Mini Kit (Qiagen).

RNASequencing libraries were generated using the SMARTer Stranded Total RNA-Seq v2 – Pico Input Mammalian kit (Takara). 100ng of RNA was used as input following the manufacturer’s protocol. Libraries were pooled together and sequenced using 75bp paired-end chemistry across 4 lanes of a Hiseq4000instrument (Illumina) to achieve 10 million reads per sample. Read quality was assessed using FastQC v.0.11.8 (Babraham Bioinformatics, UK), and SMARTer adaptors trimmed and residual rRNA reads depleted in silico using Trim galore v.0.6.4 (Babraham Bioinformatics, UK). Alignment was performed using HISAT2v.2.1.0 against the GRCh38 genome achieving a greater than 95% alignment rate. Count matrices were generated using featureCounts (Rsubreads package) and stored as a DGEList object (EdgeR package) for further analysis. Further analyses were done in R as described in (Kotagiri, Mescia et al. 2022).

### Analysis of Single-cell RNA sequencing data froms the lung adenocarcinoma cohort

#### Pre-processing, clustering, and myeloid cell type annotation

Two single-cell RNA sequencing datasets were analysed: 1) Processed Single-cell RNA sequencing UMI count data of 58 samples from 44 lung adenocarcinoma (LUAD) patients was downloaded from the NCBI Gene Expression Omnibus database (accession code <u>GSE131907</u>) ^58^. 2) Processed single-cell RNA sequencing count data of normal lung parenchyma and tumor tissues from 12 lung adenocarcinoma patients was downloaded from the Code Ocean capsule from^59^. Subsequent analyses were performed using “Seurat v3” ^60^. UMI counts were variance-stabilized using scTransform with 3000 variable features ^61^.

Top 15 principal components were used to construct UMAP embedding. Myeloid cell types were identified by scoring canonical cell type markers across clusters. Cell clustering and UMAP visualization were performed using the FindClusters and RunUMAP functions, respectively. The annotations of cell identity on each cluster were defined by the expression of known marker genes (Supplementary Table 1).

#### Functional analysis

The Monocyte-derived Macrophage (MDM) cluster was extracted for pseudobulk differential expression analysis using DESeq2 ^62^. Gene Ontology (GO) analysis was conducted using the goseq package in R ^63^ to identify enriched GO terms associated with differentially expressed genes (FDR < 0.05, LFC > 1) in tumor MDMs. A significance threshold of adjusted p-value (Benjamini-Hochberg corrected) < 0.05 was applied to determine significantly enriched GO terms. KEGG pathway enrichment analysis was conducted using pathfinder ^64^. The enrichment significance cutoff value was set to p < 0.05 for each analysis.

