## Supplementary figures for "Hypoxia drives HIF2-dependent macrophage cell cycle entry and susceptibility to lentiviral transduction" (from another version of this preprint: version 1, 2023-12-16)

### Supplementary Figure 1

A

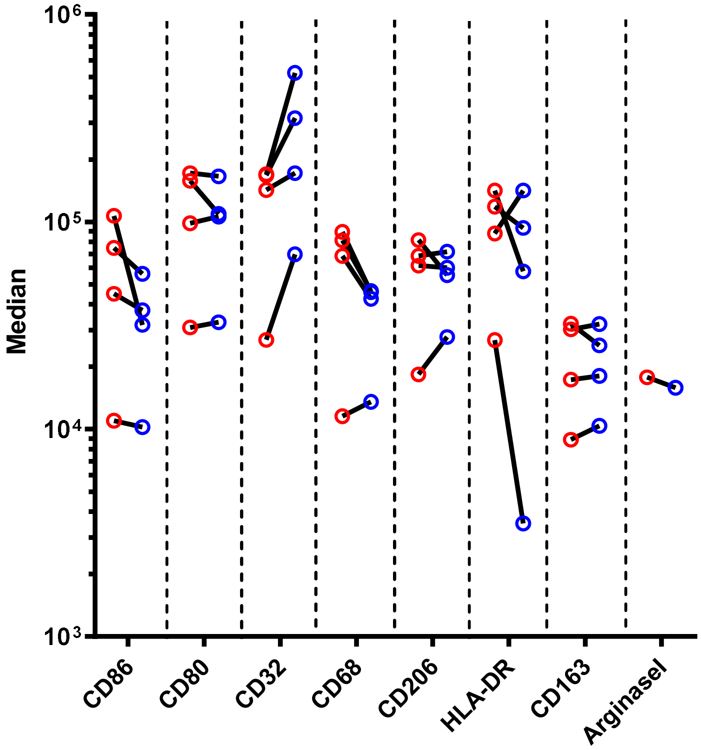


B

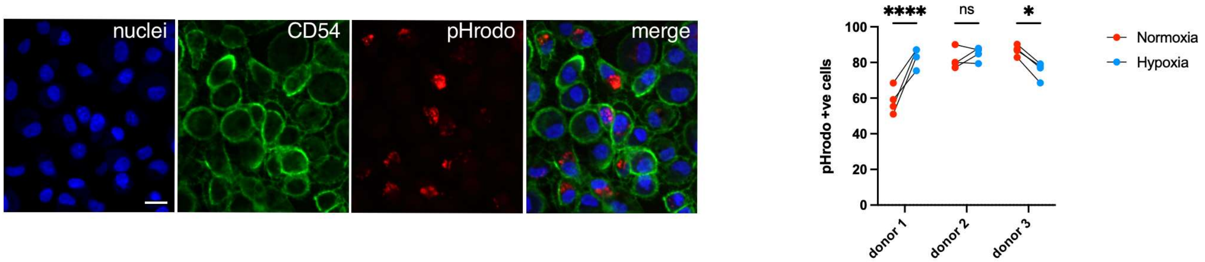

### Supplementary Figure 2

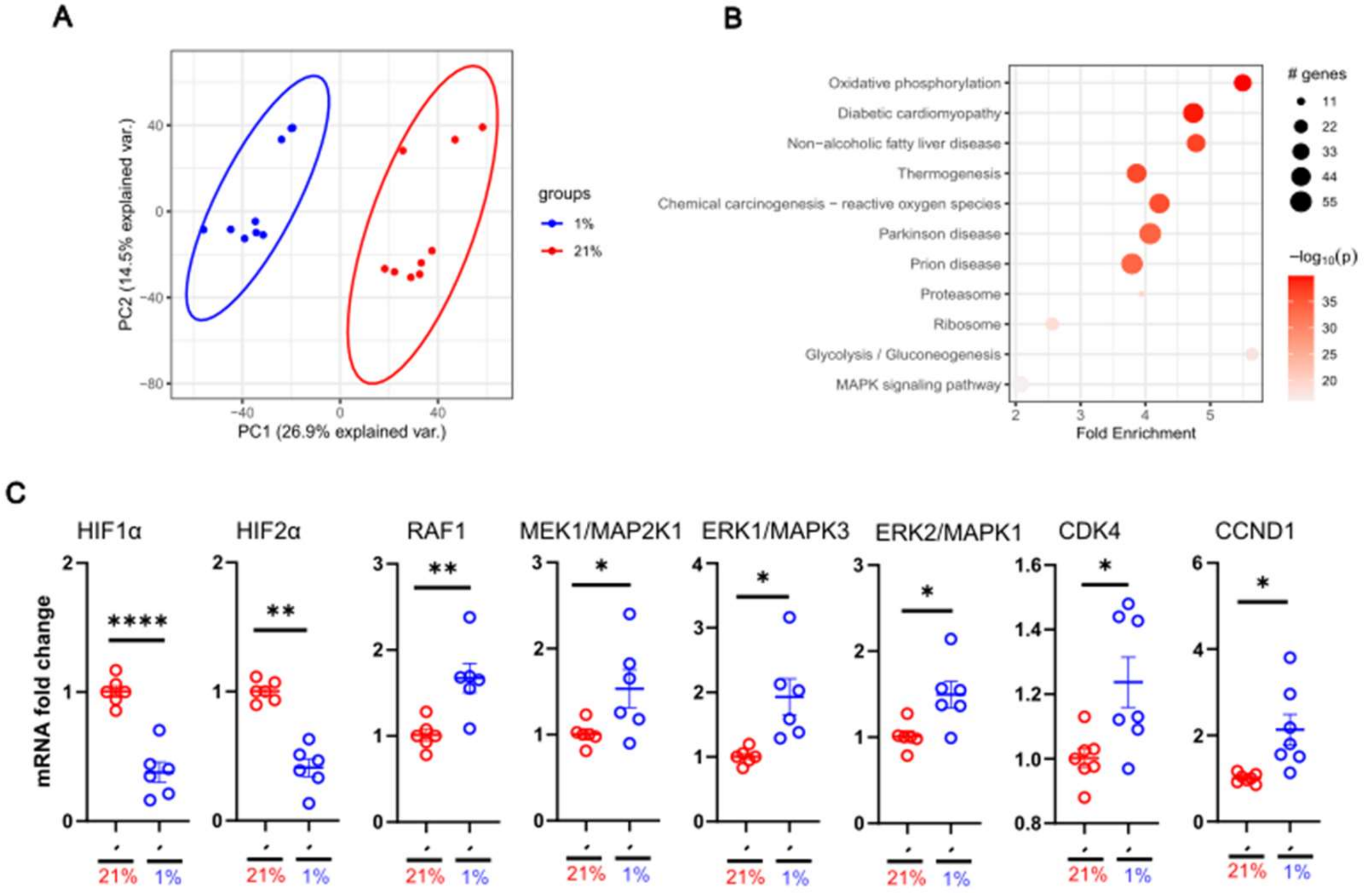

**Supplementary Figure 3**

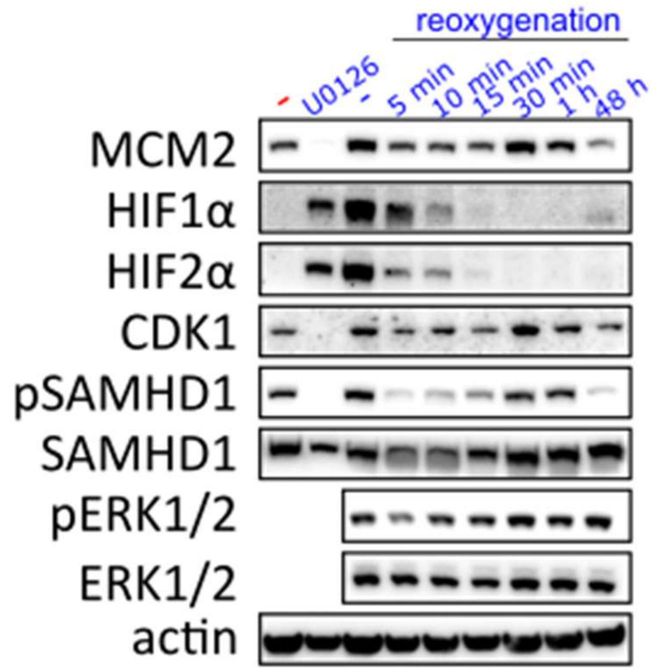

**Supplementary Figure 4**

HIV-2 infectivity

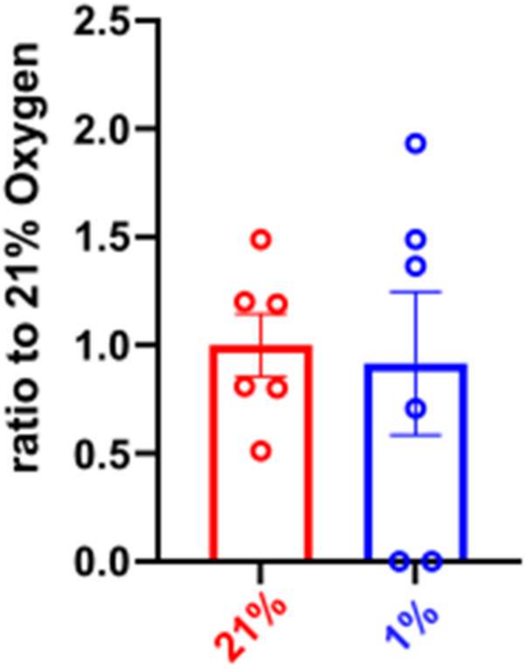

### Supplementary Figure 5

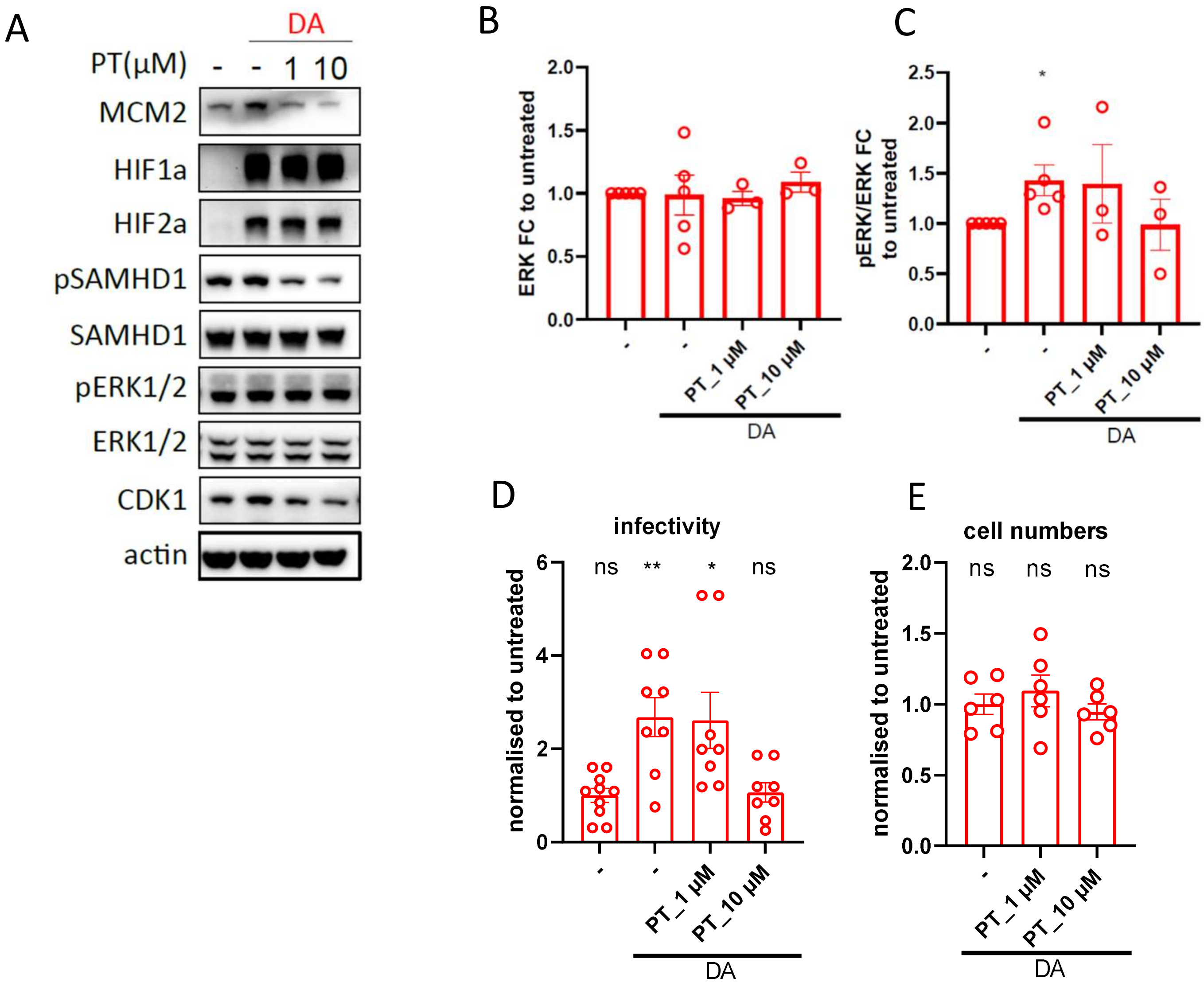

### Supplementary Figure 6

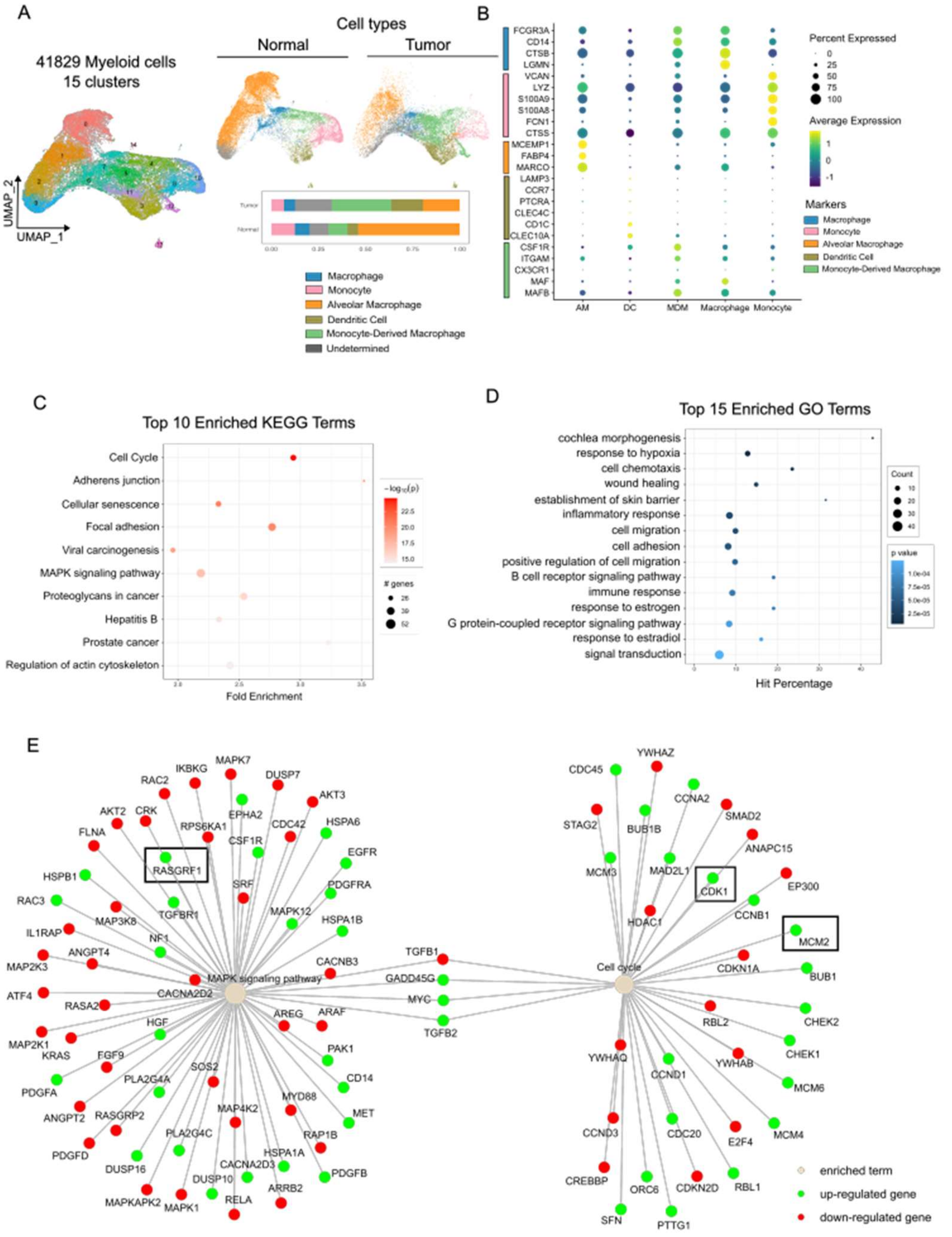
